# Physiological and molecular impairment of PV circuit homeostasis in mouse models of autism

**DOI:** 10.1101/2025.01.08.632056

**Authors:** H.R. Monday, A.M. Nieto, S.A. Yohannes, S. Luxu, K.W. Wong, F.E. Bolio, D.E. Feldman

## Abstract

Circuit dysfunction in autism may involve a failure of homeostatic plasticity. To test this, we studied parvalbumin (PV) interneurons which exhibit rapid homeostatic plasticity of intrinsic excitability following whisker deprivation in mouse somatosensory cortex. Brief deprivation reduces PV excitability by increasing Kv1 current to increase PV spike threshold. We found that PV homeostatic plasticity is disrupted in *Tsc2^+/-^*and *Fmr1^-/-^* models of autism. In wildtype mice, deprivation elevates the transcription factor ER81 which drives *Kcna1* transcription, increasing Kv1.1 protein in the axon initial segment and soma. These molecular signatures of homeostasis were absent in *Tsc2^+/-^* and *Fmr1^-/-^*. Whisker enrichment increased PV excitability, but not in *Tsc2^+/-^*, indicating that homeostasis is lost bidirectionally. Deprivation reduced feedforward L4-L2/3 inhibition in wildtype but not *Tsc2^+/-^*mice. Thus, two autism models show a convergent loss of PV circuit homeostasis at physiological and molecular levels, potentially contributing to sensory processing impairments.

## Introduction

Neurodevelopmental disorders that cause aberrant circuit function may involve a failure of homeostasis, which would otherwise compensate to stabilize cellular and circuit activity. Multiple homeostatic plasticity mechanisms coexist in cerebral cortex including homeostatic synaptic scaling of excitatory and inhibitory synapses, homeostatic intrinsic plasticity in pyramidal (PYR) cells, and homeostatic plasticity in parvalbumin (PV) inhibitory circuits^1,2^. In autism, circuit dysfunction has been proposed to involve impaired or inadequate homeostasis, which may cause circuits that are maladjusted or fragile to perturbations^3,4^. Consistent with impaired homeostasis, excitatory synaptic scaling is impaired in several mouse models of autism, including *MeCP2* ^5–7^, *Fmr1* ^8^, *Chd8* ^9^, and *Shank3* ^10,11^. Whether homeostatic plasticity in PV inhibitory circuits is impaired is unknown, but this may have outsized effects on circuit function, and possibly contribute to PV hypofunction that has been strongly linked to autism in mouse models and in post-mortem human brain tissue^12–15^.

In sensory cortex, PV interneurons receive strong feedforward and recurrent excitation, and provide perisomatic inhibition to many nearby PYR cells^16^. These PV circuits shape the sensory tuning and sensory gain of pyramidal cells. In addition, this circuit locus allows PV cells to be a critical site of firing rate homeostasis, because they are in a position to monitor local PYR activity (via recurrent excitation) and adjust their inhibitory output to homeostatically maintain PYR firing rate^13,17,18^.

In whisker somatosensory cortex (S1), PV homeostatic plasticity has been studied in response to whisker trimming or plucking, which reduces sensory drive to the deprived columns in S1^17,19,20^. A brief (1-day) deprivation of a single row of whiskers reduces the intrinsic excitability of L2/3 parvalbumin-positive (PV) interneurons in the deprived S1 columns. This results in a decrease in PV-mediated inhibition onto L2/3 pyramidal (PYR) cells, which is quantitatively balanced to offset the deprivation-induced reduction in feedforward excitation onto PYR cells. This PV plasticity is homeostatic because it stabilizes total synaptic depolarization in PYR cells, thus preserving PYR firing rate ^1,17^. Several mechanisms in PV circuits contribute to this homeostatic reduction in PV inhibition following deprivation, but the most rapid component is a decrease in PV intrinsic excitability, which occurs at 1 day of deprivation, prior to detectable weakening of excitatory synapse onto PV cells^20^. Decreased PV excitability is associated with an increase in voltage-activated delayed rectifier (likely Kv1) currents in PV cells, which regulate near-threshold PV excitability and spike threshold^21^. The molecular pathways that mediate this plasticity are unknown, but could involve upregulation of *Kcna1* expression (the gene for Kv1.1) by the activity-dependent transcription factor ER81(also called ETV1) ^22^, which has been shown to modulate ion channel expression and excitability in PV cells and other interneurons in response to strong genetic or pharmacological reductions of network activity^23,24^. Whether the ER81-Kv1.1 pathway is engaged by natural changes in sensory activity on rapid timescales is unknown.

Here, we test the hypothesis that PV homeostasis is impaired in autism spectrum disorder (ASD) by examining two genetically distinct ASD mouse models *(Fmr1^-/-^* and *Tsc2^+/-^*). Both models exhibit hypofunctional feedforward inhibition in S1^12,25^, but whether these or other autism models exhibit aberrant PV homeostasis is unknown. Loss-of-function mutation in *Fmr1* causes Fragile X Syndrome (FXS), the most common monogenic cause of ASD. *Fmr1* encodes Fragile X Messenger Ribonucleoprotein (FMRP), an RNA-binding protein whose loss is linked to dysregulated activity dependent protein synthesis^26,27^ and, typically, an overall increase in protein synthesis^28–31^, though this is somewhat controversial^32^. Tuberous sclerosis complex (TSC) is another syndrome associated with ASD that results from mutation in *Tsc1* or *Tsc2* genes that leads to neurological symptoms including epilepsy, autism, cognitive disability and neurobehavioral abnormalities^33,34^. Because TSC2 acts as a brake on mTOR activation, *Tsc2* mutation was initially thought to cause exaggerated protein synthesis, similar to *Fmr1* mice^35^, but more recent evidence suggests reduced protein synthesis^36,37^. Because *Tsc2^+/-^* and Fmr1*^-/-^* mice differ at the synaptic, biochemical, and cognitive levels^25^, they provide a strong test for potential convergence in disruption in PV homeostasis.

We found that *Tsc2^+/-^* and *Fmr1^-/-^* mice share a profound impairment in PV circuit homeostasis, in which 1 day of whisker deprivation completely fails to reduce PV excitability in either mouse model. On the molecular level, PV homeostasis in wildtype (WT) mice was accompanied by deprivation-induced upregulation of ER81, rapid transcription of *Kcna1* mRNA, and increased Kv1.1 protein both in bulk in L2/3 PV soma and at the AIS. ER81 and Kv1.1 regulation by deprivation were entirely absent in *Tsc2^+/-^* and *Fmr1^-/-^*. We found that the combination of sparing one row of whiskers plus environmental enrichment drove an opposing increase in PV excitability and decreased Kv1.1 expression, indicating the PV homeostasis is bidirectional. Both directions of homeostatic plasticity were lost in *Tsc2^+/-^*mice. At the circuit level, *Tsc2^+/-^*mice lacked deprivation-induced weakening of L4-L2/3 inhibition. Thus, these models exhibit a convergent deficit in rapid PV circuit homeostasis, which may contribute to sensory processing impairments.

## Results

### Homeostatic PV intrinsic plasticity is impaired in *Tsc2^+/-^* and *Fmr1*^-/-^ mice

To assay PV intrinsic plasticity, we labeled PV cells using an enhancer virus (AAV-S5E2-dTom- nlsdTom) injected into S1 of *C57* wildtype (*WT-C57*), *Tsc2^+/-^*, and *Fmr1-/-* mice at postnatal day (P) 3^38^. Between P18-P23, mice underwent D-row whisker plucking or sham-plucking, followed by whole-cell recording from PV cells 24 hours later (**Figure 1A-B**). The virus was ∼80% specific and sensitive for PV cells when injected at P3, based on immunostaining with anti-PV antibody (**Figure 1C**). We made in vitro sections of S1 in the ’across-row’ plane that allows identification of specific whisker-related columns, and recorded from tdTom+ cells in L2/3 of the deprived D column. tdTom+ cells that exhibited fast-spiking (FS) spike patterns were considered PV cells (**Figure 1B,D**). To assess PV intrinsic excitability, we recorded in current clamp, identified rheobase and then recorded F-I curves at current levels above rheobase. In *WT-C57* mice, F-I curves were significantly reduced in 1-day deprived mice relative to sham mice with intact whiskers (**Figure 1D,E**), associated with an increase in spike threshold (**Figure 1F,H**) and in the latency to first spike (**Figure 1G**). Passive properties (Vrest and Rinput), rheobase, and spike shape were not affected (**Figure S1**). These findings in *WT-C57* mice replicate the reduction in L2/3 PV intrinsic excitability measured from genetically identified PV cells in PV-Cre;tdTomato mice ^20^.

**Figure 1.**
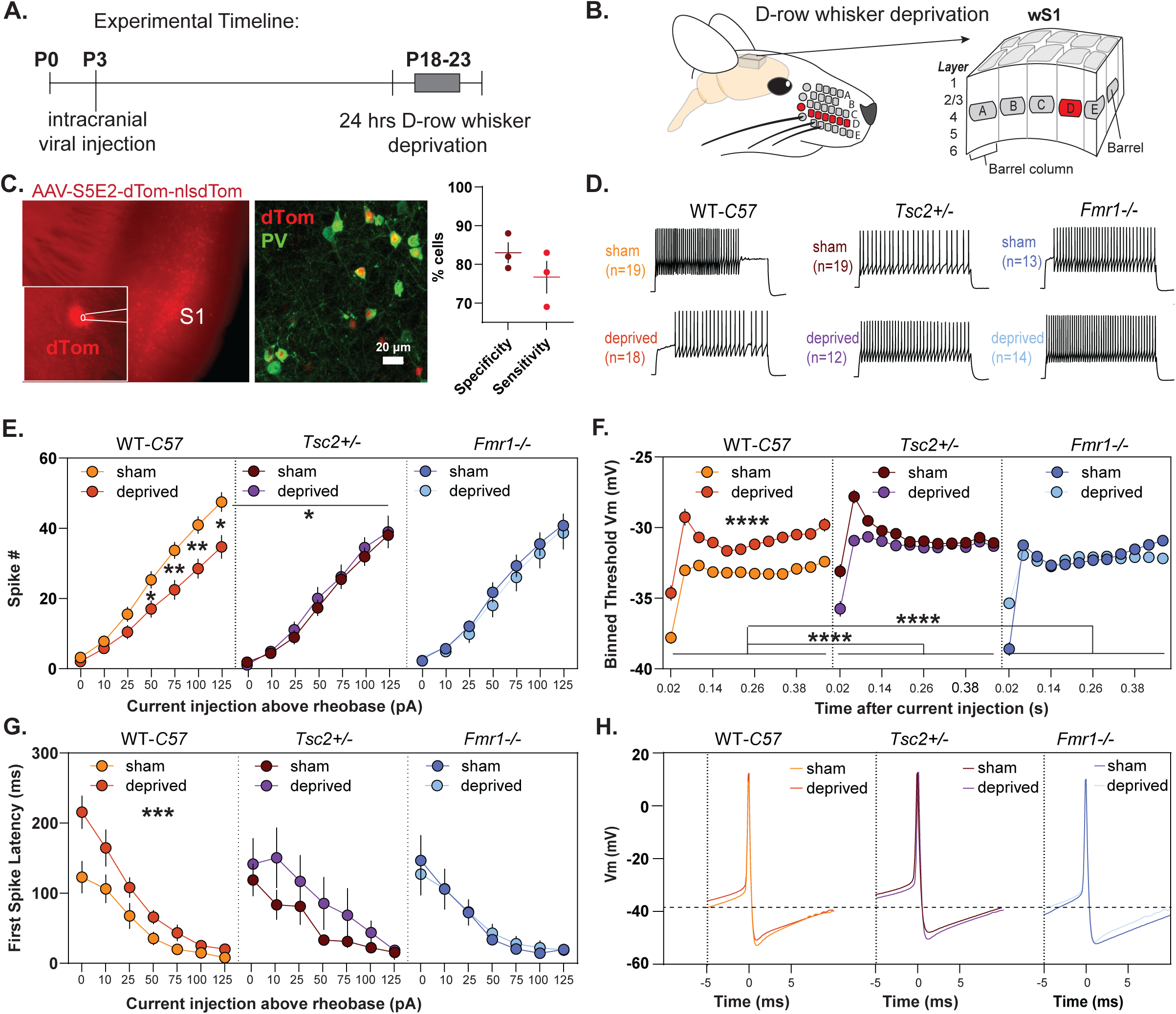
Deprivation-induced weakening of PV intrinsic excitability is absent in Tsc2+/- and Fmr1-/- mice. A. Timeline of PV intrinsic excitability recording experiments B. Whiskers in the D-row are plucked under anesthesia and mice are returned to their cage for 24 hours. For sham conditions, mice are anesthetized, and whiskers are gently stroked. To identify and record from the deprived column, ‘across-row’ sections (see Methods) of S1 are made that contain one barrel from each row. C. (left) Representative image of intracranial viral injection targeting S1. (Inset) Image of patch pipette on confirmed fluorescent PV cell. Colocalization of PV immunostaining and dTom viral expression revealed 83% specificity (# dTom+, PV+ cells/ all dTom+ cells) and 78% sensitivity (# PV+, dTom+ cells/ all PV+ cells), n=3 mice. D. Representative traces of current injections from L2/3 PV interneurons at 100 pA above rheobase in WT-C57, Tsc2+/-, and Fmr1-/- sham and deprived mice E. Whisker deprivation reduced PV spiking in WT-C57 mice, while ASD models did not change their firing. WT-C57 sham mice have significantly increased firing compared to Tsc2 sham. Two-way RM-ANOVA with post-hoc Dunnett’s test, *p<0.05, **p<0.01. n = 19, 20, 19, 12, 13, 14 cells respectively from 5-7 mice per group. WT sham v. *Tsc2* sham curves were significantly different (*p < 0.05) but WT sham v. *Fmr1* sham were not. F. Whisker deprivation increased the spike threshold in WT-C57, but not Tsc2+/- and Fmr1 -/-. Threshold was increased in both autism models compared to WT sham. Threshold was measured in all spikes at 0, 10, 25, and 50 pA above rheobase. Two-way RM-ANOVA with post-hoc Dunnett’s test, n = 3843, 2473, 2901, 2086, 2424,1800 spikes respectively G. The latency to the first spike was significantly increased by whisker deprivation in the WT-C57, but not in either autism model. Two-way RM-ANOVA with post-hoc Dunnett’s test, ***p<0.001. n = 19, 20, 19, 12, 13, 14 cells respectively from 5-7 mice per group. H. Average spike waveforms from each condition reveal changes in spike threshold.

Unlike in WT mice, deprivation caused no change in F-I curves, spike threshold, or spike latency in either *Tsc2^+/-^* or *Fmr1^-/-^* mice (**Figure 1D-G**), and no change in passive properties or spike shape (**Fig. S1**). F-I curves in sham mice were weaker in *Tsc2^+/-^*than *WT-C57* (**Figure 1D,E**), consistent with (^39^), and PV spike threshold in sham mice was elevated in*Tsc2^+/-^* and *Fmr1^-/-^* compared to *WT- C57* (**Figure 1F,H**). These findings indicate that PV hypofunction that has been observed under baseline sensory conditions in *Tsc2^+/-^* and *Fmr1^-/-^* mice ^40^ involves, in part, abnormally weak PV intrinsic excitability. Vrest, Rinput, rheobase, and spike shape were not significantly different across genotypes, except for subtly broader spike width in *Tsc2^+/-^* PV cells (**Figure S1**). Thus, homeostatic reduction of PV intrinsic excitability is absent in these autism models.

### Deprivation alters Kv1.1 expression and length of the axon initial segment

In PV cells, Kv1.1 channel insertion in the axon initial segment (AIS) elevates PV spike threshold and delays spike onset^21^. To assess whether the spike threshold and latency effects of whisker deprivation during PV circuit homeostasis reflect increased Kv1.1 at the AIS, we performed immunostaining on S1 sections from sham and deprived *WT-C57* mice. Cortical columns corresponding to D-row whiskers were identified in across-row sections using PV and vGlut2 staining (**Figure 2A**). AnkG antibody was used to mark AIS compartments and Kv1.1 antibody to measure Kv1.1 protein levels. We located and imaged the AIS of individual L2/3 PV cells (labeled with PV antibody) in D columns with high-resolution Airyscan confocal imaging. Kv1.1 staining intensity was measured within the PV AIS (**Figure 2B,C,D**). Imaging was performed blind to condition. Data was averaged across approximately 10-20 PV cells per column. Kv1.1 staining in the AIS was increased in mice after 1-day deprivation compared to sham mice (**Figure 2E**). Deprivation also decreased PV AIS length, a finding that is associated with reduced excitability in multiple cell types ^41,42^ (**Figure 2F**). The distance of the AIS from the cell soma—another correlate of cellular excitability—was not altered (**Figure 2G**). Thus, deprivation increases Kv1.1 expression in the AIS, and reduces AIS length in PV cells during homeostasis.

**Figure 2.**
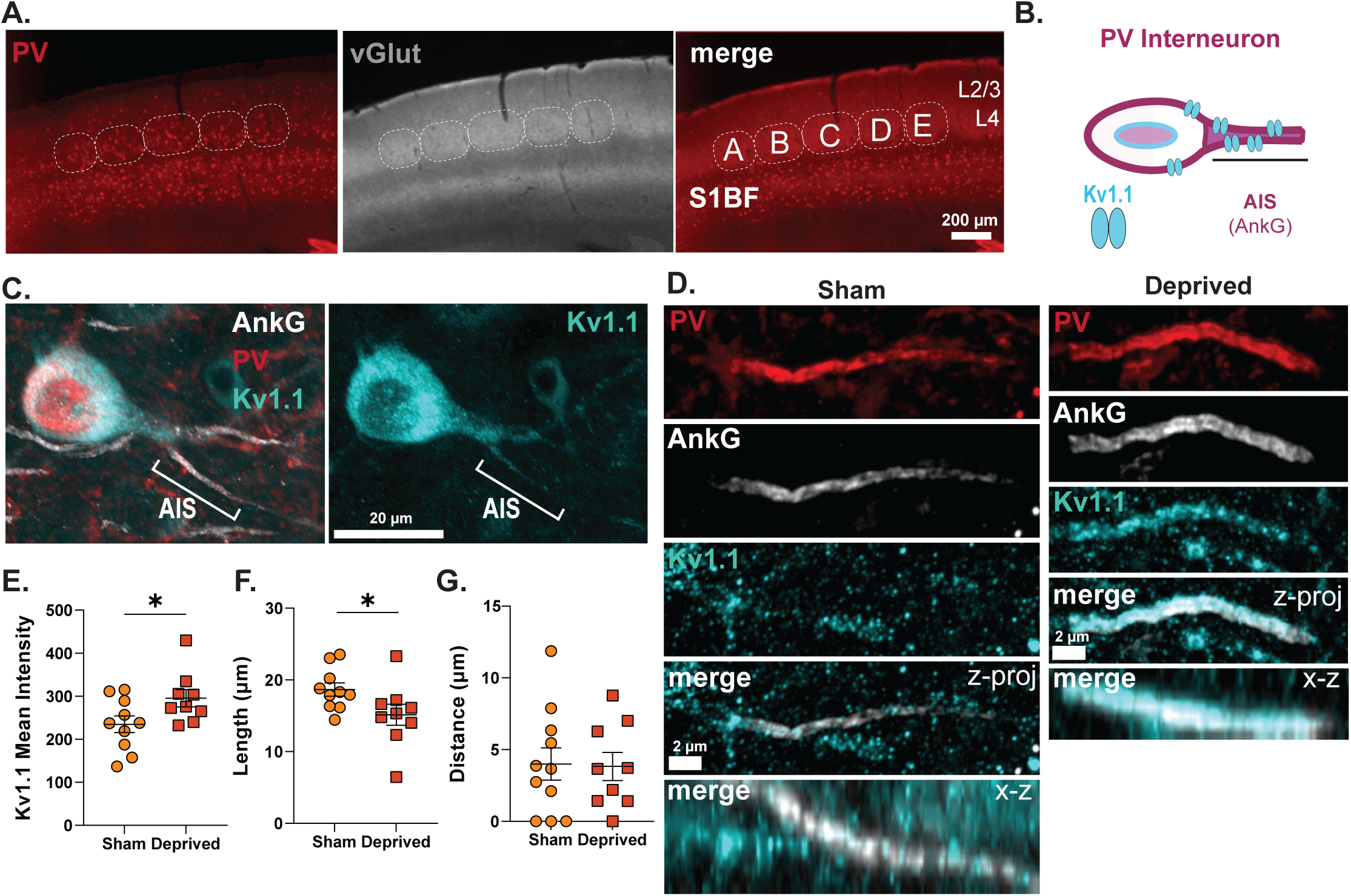
Deprivation triggers plasticity of the PV axon initial segment. A. Representative widefield 5X image of PV and vGlut staining in S1. Note barrels visible in L4. B. Schematic of AIS Kv1.1 channel expression in the AIS of PV interneurons C. Representative 63X airyscan confocal image of AnkG immunolabeled AIS in PV interneuron expressing Kv1.1 D. Representative magnified maximum intensity projections showing Kv1.1 expression in AnkG labeled axons from Sham and Deprived mice. E. Mean Kv1.1 staining intensity is increased in AIS’s from deprived mice. Nested t-test, p = 0.04 F. AIS length is decreased in deprived mice. Nested t-test, p = 0.04 G. The distance from the soma at which the AIS begins is unchanged in sham v. deprived mice. Nested t-test, p = 0.85 a. (n = 10, 9 slices 5 mice each)

### Whisker deprivation increases Kv1.1 and ER81 expression in PV cells

Typically, activity-dependent gene expression is triggered by increases in activity, whereas whisker deprivation drives an acute, modest reduction in L2/3 network activity ^43^. A previous study identified a molecular pathway by which decreased PV activity increases transcription factor ER81 levels in PV cells, which binds to the *Kcna1* promoter (the gene for Kv1.1) and drives increased Kv1.1 expression^23^. However, it is unclear whether this pathway is recruited by natural changes in sensory activity at rapid time scales or is involved in autism pathophysiology. We hypothesized that whisker deprivation-induced changes in network activity would lead to increased ER81, and this could promote Kv1.1 expression (**Figure 3A**). To test whether bulk Kv1.1 expression or ER81 was altered by deprivation in PV cells, we analyzed Kv1.1 and ER81 staining in L2/3 PV somata in D whisker columns (**Figure 3B**). In L2/3, Kv1.1 staining was mainly observed in PV cells. In *WT-C57* mice, deprivation increased Kv1.1 staining intensity in L2/3 PV cells by ∼ 28% (**Fig. 3B-D**). This increase was entirely absent in *Tsc2^+/-^* and *Fmr1*-/- mice. Among sham mice, *Tsc2^+/-^* had similar Kv1.1 expression as WT, but *Fmr1^-/-^* had substantially reduced Kv1.1, suggesting that other conductances compensate to maintain PV excitability (**Figure 3C-D**).

**Figure 3.**
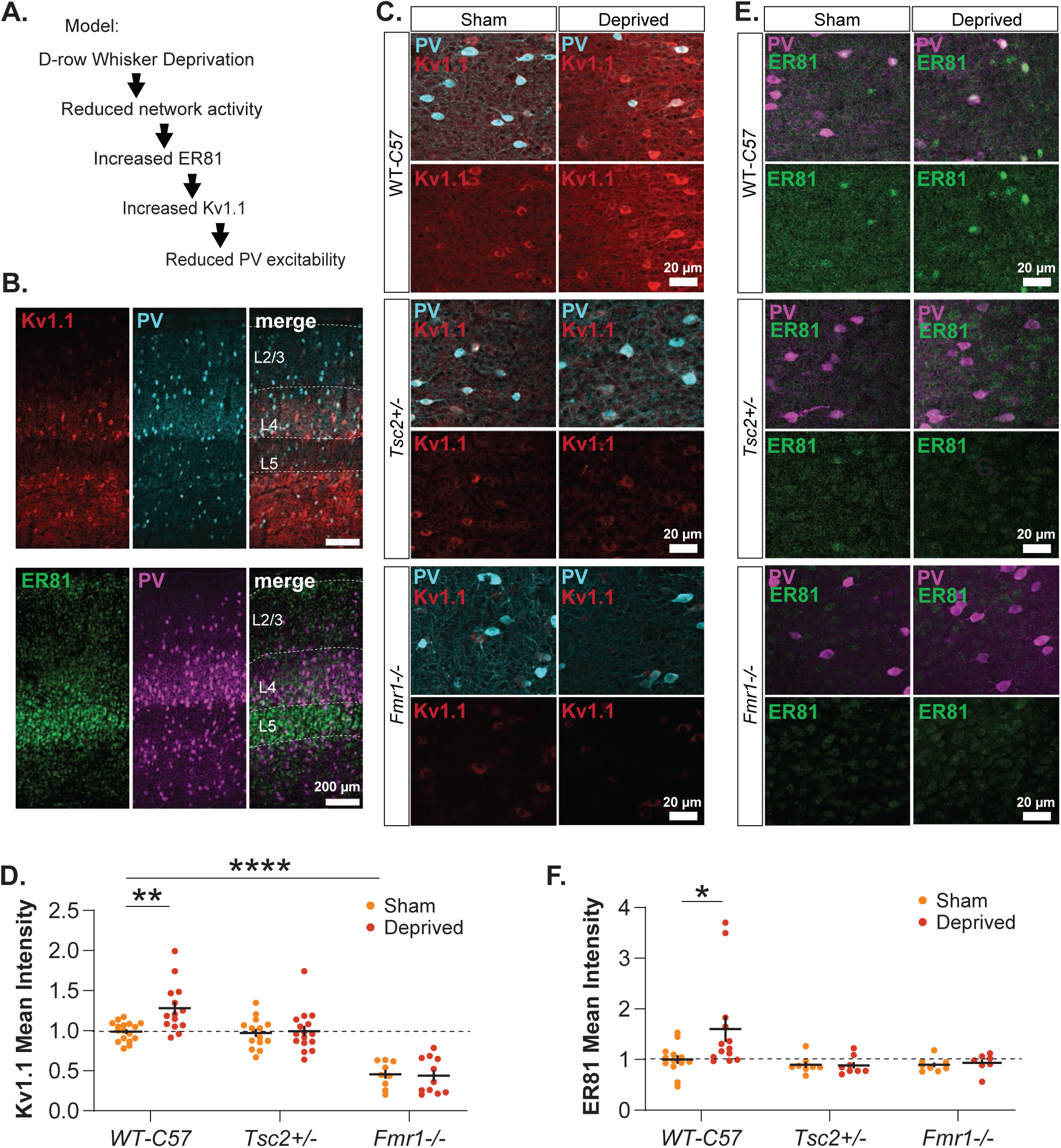
Whisker deprivation increases Kv1.1 and ER81 expression in PV cells. A. Hypothesized model of the molecular mechanism of PV circuit plasticity: Whisker deprivation triggers reduction in feedforward activity which increases transcription factor ER81 levels and promotes Kv1.1 expression. B. Widefield images of Kv1.1 (red) and PV (cyan) immunostaining (top) and ER81 (green) and PV (magenta) (bottom). Kv1.1 is primarily expressed in PV cells. ER81 is highly expressed in L5 pyramidal cells. C. Representative confocal images of Kv1.1 (red) and PV (cyan) immunostaining in D column of Layer 2/3 D. Representative confocal images of ER81 (green) and PV (magenta) immunostaining in D column of Layer 2/3 E. Mean Kv1.1 staining intensity inside L2/3 PV cells is increased with whisker deprivation in *WT-C57* but not autism mouse models. Kv1.1 levels are significantly reduced at baseline in *Fmr1-/-* mice. Two-way ANOVA with Tukey’s test for Multiple Comparisons, Genotype: F(2,75) = 62.10, p <0.0001, n = 17,14,14,16,9,11 slices respectively, 4-6 mice per group F. Mean ER81 staining intensity inside L2/3 PV cells is increased with whisker deprivation in *WT-C57* but not autism mouse models. Two-way ANOVA with Tukey’s test for Multiple Comparisons, Genotype: F(2,51) = 4.757, p = 0.013, n = 13,13,8,8,8,7 slices respectively, 4-6 mice per group

In *WT-C57* mice, ER81 increased following 1 day of whisker deprivation, but this effect was absent in *Tsc2^+/-^* and *Fmr1^-/-^* mice (**Figure 3E,F**). Among sham mice, *Tsc2^+/-^*and *Fmr1^-/-^*showed a nonsignificant trend towards reduction of ER81 relative to WT (**Figure 3E,F**). These findings are consistent with the hypothesis that whisker deprivation triggers increased production of the transcription factor ER81, which promotes Kv1.1 expression, thus reducing PV intrinsic excitability. The absence of activity-dependent regulation of ER81 and Kv1.1 could potentially explain the loss of homeostatic PV intrinsic plasticity in these two mouse models.

### Kcna1 is rapidly transcribed following whisker deprivation

To directly test whether deprivation regulated the transcription of *Kcna1*, we measured *Kcna1* mRNA in L2/3 PV cells. We used quantitative HCR-FISH for *Kcna1* mRNA, coupled with the big-FISH software pipeline for analysis (**Figure 4A,B**). *PV* mRNA was used to identify PV cells and cells were manually outlined to create ROIs. Surprisingly, we observed a trend towards reduced *Kcna1* at 24 hours after whisker deprivation (**Figure 4C-E, H**). Given that Kv1.1 protein levels is already upregulated at 24 hours, we hypothesized that *Kcna1* mRNA is upregulated on a faster timescale, followed by mRNA degradation. Repeating the HCR-FISH at 1.5 and 5 hours of deprivation, we observed that Kcna1 levels were significantly upregulated at 1.5 hours, and that this declined to normal levels at 5 hours (**Figure 4C-G**). This was true in both the somatic, nuclear, and cytosol compartments (**Figure 4C-F, Figure S4**). Thus, *Kcna1* transcription is rapidly upregulated by whisker deprivation, within 1.5 hours. This demonstrates that PV cells detect altered whisker experience and launch the homeostatic response much more rapidly than had been previously known.

**Figure 4.**
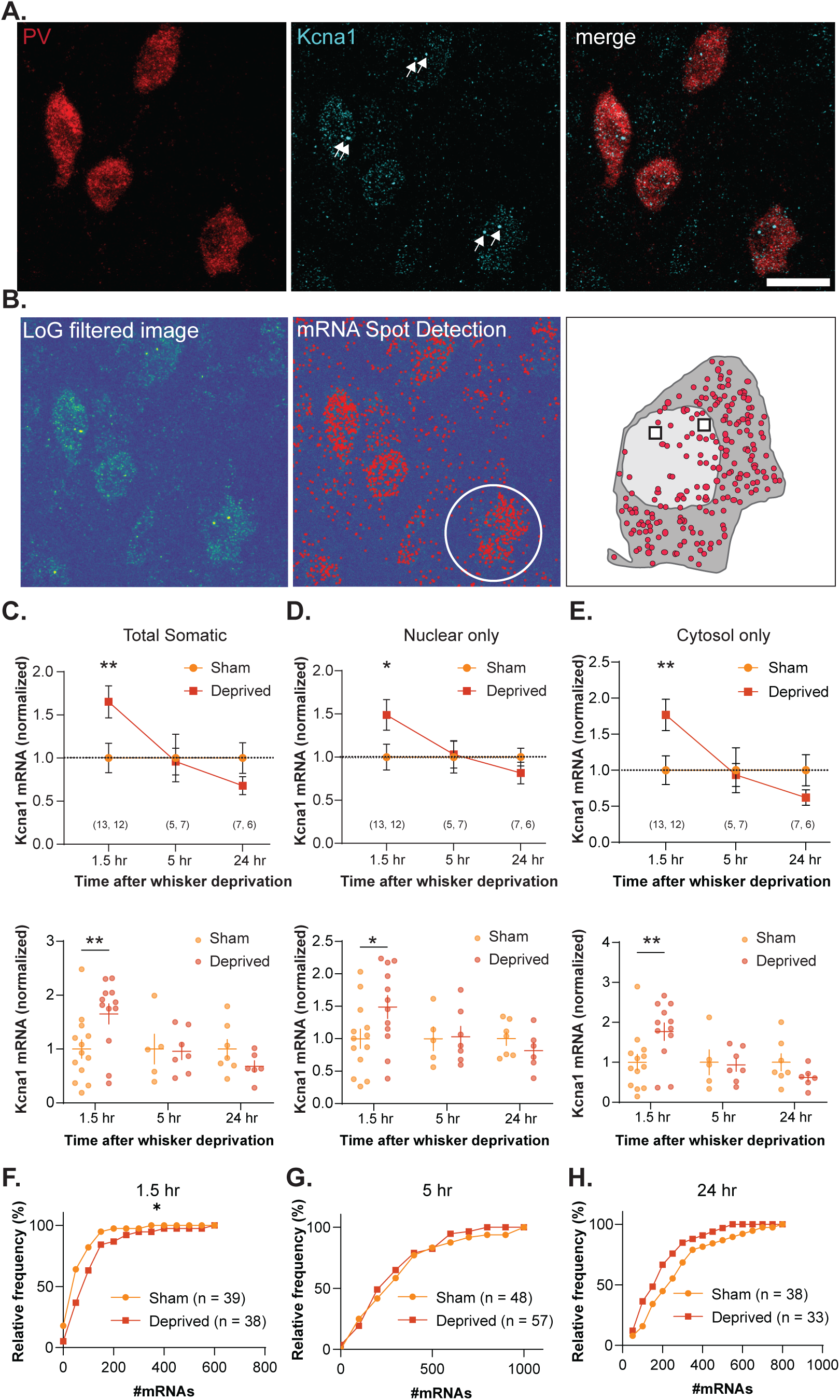
Kcna1 is rapidly transcribed following whisker deprivation in WT mice. A. Representative confocal images of HCR-FISH. Kcna1 is highly expressed in PV cells. White arrows indicate putative transcription sites. B. Example of BIG-FISH analysis pipeline. Images are pre-processed with a LoG filter before spot detection. Spots are de-clustered and localized to segmented cells. C. Total somatic Kcna1 mRNA inside PV cells is significantly increased at 1.5 hour following whisker deprivation but back to baseline by 5 hours and slightly reduced at 24 hours. Two Way ANOVA with Tukey’s test for Multiple Comparisons, ** p < 0.01, n = slices, 1-7 PV cells are measured per slice, 3-6 mice per group. (top) Summary plots are normalized to sham condition. Symbols and lines represent mean + SEM of slices. (bottom) Scatterplot showing all points (slices), lines represent mean + SEM of slices. D. Nuclear Kcna1 mRNA is significantly increased at 1.5 hours post-whisker deprivation. Two Way ANOVA with Tukey’s test for Multiple Comparisons. * p < 0.05. Summary plots are normalized to sham condition. Points and lines represent mean + SEM of slices. 1-20 PV cells are measured per slice. n = (slices, animals) E. Cytosolic Kcna1 mRNA is significantly increased at 1.5 hours post-whisker deprivation. Two Way ANOVA with Tukey’s test for Multiple Comparisons. ** p < 0.01. Summary plots are normalized to sham condition. Points and lines represent mean + SEM of slices. 1-20 PV cells are measured per slice. n = (slices, animals) F. Deprivation upshifts the distribution of # mRNAs per cell at 1.5 hours. Kolmogorov-Smirnov test, p < 0.05, n = cells G. No change in the distribution of mRNAs per cell at 5 hours. Kolmogorov-Smirnov test, p = 0.05, n = cells H. No significant change in the distribution of mRNAs per cell at 24 hours after deprivation, but a trend towards reduction. Kolmogorov-Smirnov test, p = 0.154, n = cells

### Homeostatic PV intrinsic plasticity is bidirectionally impaired in *Tsc2^+/-^* mice

If PV intrinsic plasticity indeed functions as a rapid homeostat to stabilize pyramidal cell firing, then it should be bidirectional, i.e. reducing sensory input should reduce PV excitability (as shown above and previously^20^), and enhancing sensory input should increase PV excitability. To test this latter prediction, we devised a ‘D-sparing plus enrichment’ protocol wherein the D row of whiskers are spared (by plucking all other contralateral whisker rows, and all ipsilateral whiskers, which is the opposite of our deprivation protocol) and mice are placed in an enriched environment with textured objects and additional littermates, to encourage use of the remaining D-row whiskers (**Figure 5A-B**). After injecting with the S5E2 enhancer virus at P3, littermates at P18-23 were randomly assigned to the ’sparing plus enrichment’ group or the ’standard experience’ group that underwent sham plucking and were housed in the standard cage environment. In separate experiments, we confirmed that the ’D-sparing plus enrichment’ protocol drove activity in L2/3 of the D whisker columns, using immunostaining for c-fos, an immediate early gene whose expression is highly correlated with cellular activity. C-fos was strongly induced in a D column-specific manner 2 hours after whisker enrichment (**Figure S5A**). To test for PV intrinsic plasticity, we made whole-cell recordings from L2/3 PV cells in D columns after 1-3 days, using the same methods in Figure 1. In *WT-C57* mice, D- sparing plus enrichment significantly increased PV excitability measured by F-I curves, relative to littermates in the standard experience group (**Figure 5C-D**). Unlike with whisker deprivation, D- sparing plus enrichment in *WT-C57* mice did not alter spike threshold or spike latency, potentially indicating a different mechanism of expression (**Figure 5E-G**). Passive properties and spike shape were not altered (**Figure S5B-G**).

**Figure 5.**
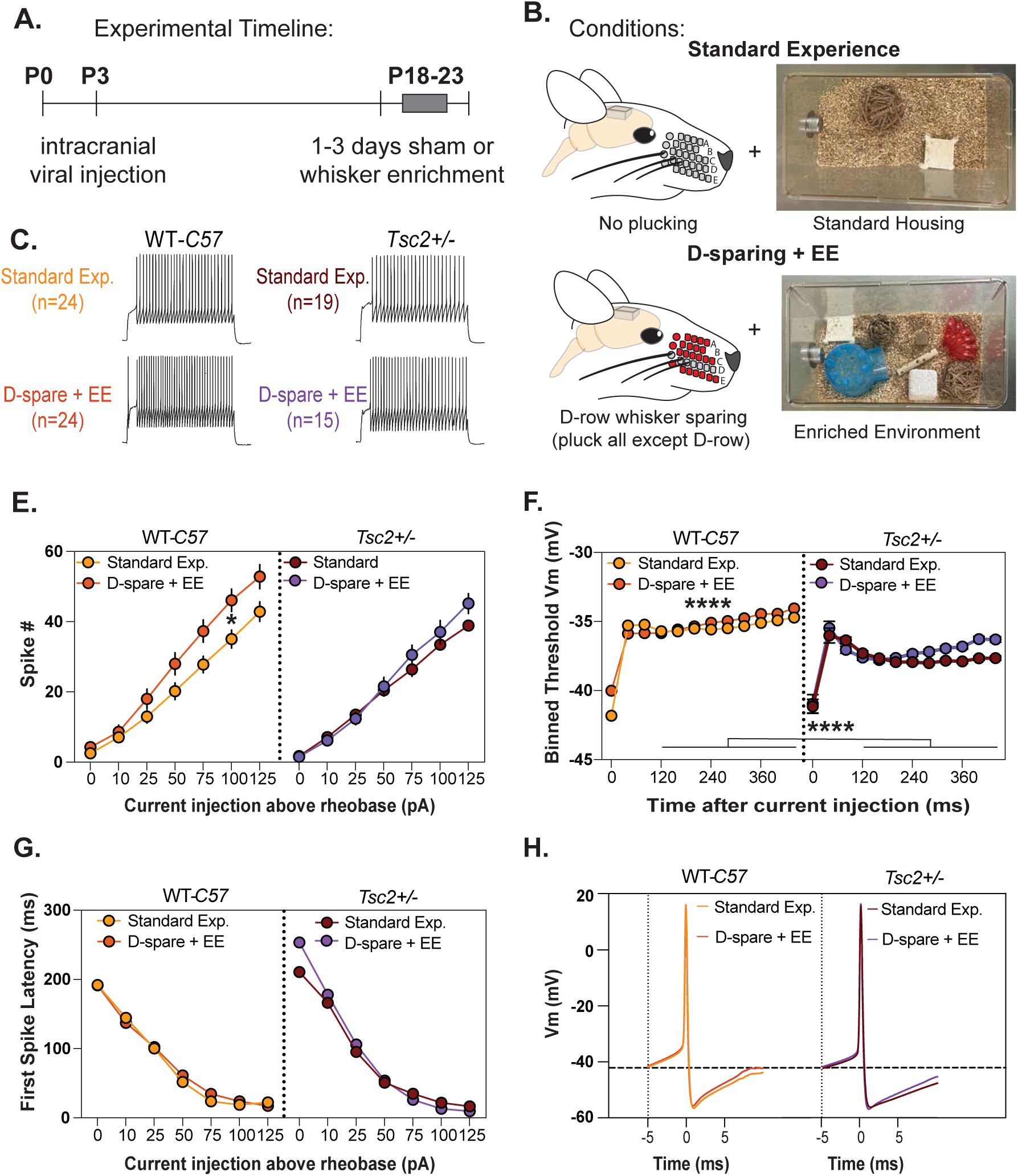
Enrichment plus sparing increases PV intrinsic excitability in wild type but not Tsc2+/- mice. A. Timeline of PV intrinsic excitability recording experiments. B. Experience conditions for whisker enrichment: ‘Normal experience’ mice are anesthetized and whiskers are gently stroked,then weaned into standard housing cages for 1-3 days. Standard housing consisted of 2 mice per cage. (right) Representative image of enriched environment. ‘Sparing + enrichment’ mice underwent plucking of all whiskers except the D-row on the right side of the face (D-sparing). After plucking, whisker D-spared mice are weaned into enriched environment cages for 1-3 days. Enriched environment consisted of 3 mice per cage and included a plastic hut, plastic tube, and a mixture of sticks, blocks, and spheres of various textures. C. Representative traces of current injections from L2/3 PV interneurons at 75pA above rheobase in WT-C57 and Tsc2+/- normal experience and sparing + enrichment mice. D. Sparing + enrichment increases PV spiking in WT-C57, but not Tsc2+/- mice. Two-way RM-ANOVA with post-hoc Dunnett’s test, *p<0.05. WT-C57 normal experience: n=24 cells from 7 mice; WT-C57 sparing + enrichment: n=24 cells from 6 mice; Tsc2+/- normal experience: n=19 cells from 5 mice; Tsc2+/- sparing + enrichment: n=15 cells from 5 mice. E. Spike threshold is significantly altered by genotype and experience compared to WT-C57 mice. Threshold was measured in all spikes at 0, 10, 25, and 50 pA above rheobase. Two-way RM-ANOVA with post-hoc Dunnett’s test. WT-C57 normal experience: n=4,723 spikes; WT-C57 sparing + enrichment: n=6,553 spikes; Tsc2+/- normal experience: n=3,756 spikes; Tsc2+/- sparing + enrichment: n=2,904 spikes. F. Sparing + enrichment has no effect on the latency to the first spike in WT-C57 or Tsc2+/- mice. Two-way RM-ANOVA with post-hoc Dunnett’s test. G. Averaged PV action potential waveform for WT-C57 and Tsc2+/- normal experience and sparing + enrichment mice. Average of all action potentials, excluding the first spike of the train, at 0, 10, 25, and 50pA above rheobase.

To test if PV intrinsic plasticity was bidirectionally impaired in autism, we focused on *Tsc2^+/-^* mice. In the standard experience condition, *Tsc2^+/-^* mice had weaker F-I curves than in WT mice (**Figure 5C-D**), matching the results from Figure 1. In *Tsc2^+/-^* mice, sparing plus enrichment did not increase F-I curves relative to standard experience, indicating a loss of upward PV intrinsic plasticity (**Figure 5C-F**), again with no changes in passive properties or spike shape (**Figure S5B-G**). Thus, experience-dependent plasticity of PV intrinsic excitability is bidirectional, as presumed of a homeostatic mechanism, and both directions of homeostatic plasticity are lost in *Tsc2^+/-^* mice.

### Homeostatic plasticity of feedforward inhibition and excitation is disrupted in *Tsc2^+/-^*

To understand the consequences of loss of PV intrinsic plasticity on circuit function, we assessed feedforward excitation and inhibition in the L4-L2/3 circuit (**Figure 6A**). L4-L2/3 feedforward inhibition is primarily mediated by PV cells ^19,44,45^. In S1 slices, we stimulated in L4 of the D column, recorded from a L2/3 PYR cell in the same column, and measured feedforward EPSCs and IPSCs by holding the PYR cell at -68 mV or 0 mV (E_Cl_ or E_AMPA_). We first identified the smallest L4 stimulation intensity that evoked a detectable EPSC, defined as E-threshold (Eθ). We then recorded at 1.0, 1.2 and 1.4 x Eθ to sample the input-output curve. In these conditions, 85% of the evoked IPSC is due to disynaptic feedforward inhibition ^20^. In *WT-C57* mice, 1 day of whisker deprivation significantly reduced feedforward inhibition and more modestly reduced feedforward excitation relative to sham littermates (**Figure 6B-D**). The reduction in feedforward inhibition is known to reflect deprivation-induced weakening of PV intrinsic excitability ^20^.

**Figure 6.**
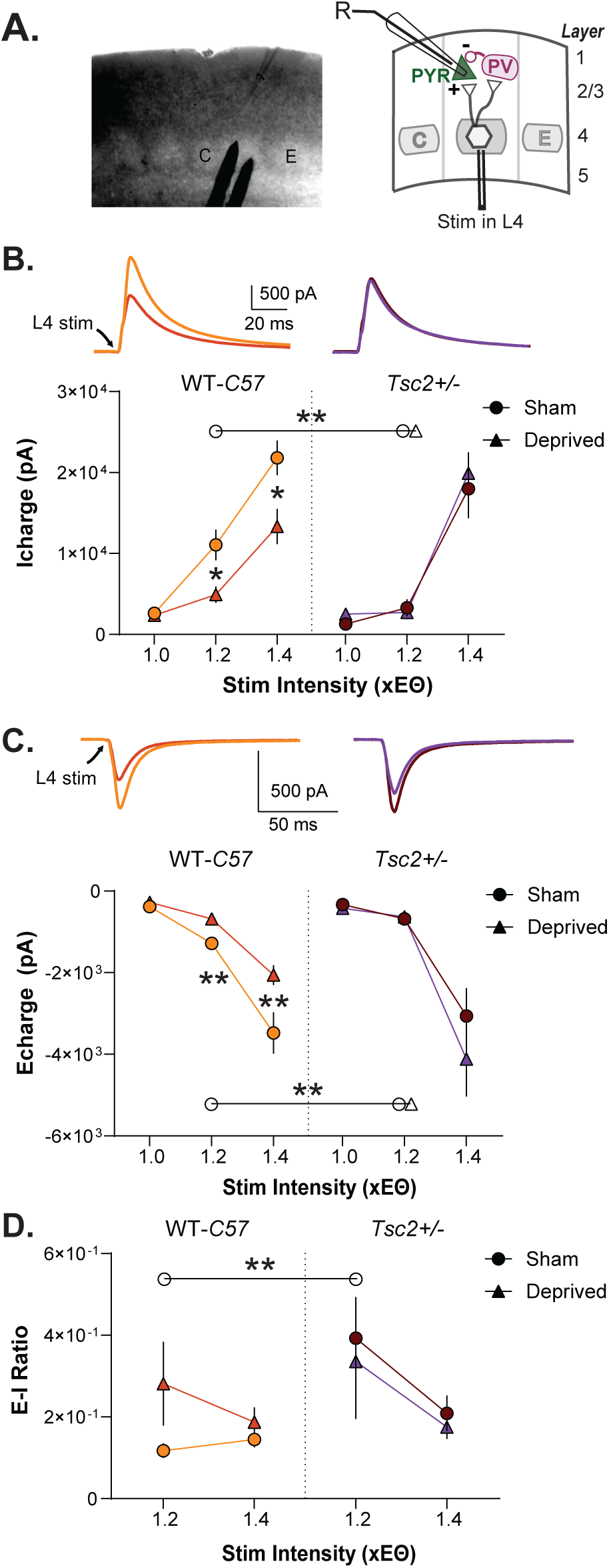
Homeostatic plasticity of feedforward inhibition and excitation is impaired in the Tsc2+/- mouse model. A. Image (left) and schematic (right) of voltage clamp recording configuration in L2/3 B. Deprivation weakens L4-L2/3 feedforward inhibition in wildtype but not Tsc2+/- mice. Tsc2+/- mice show weakened inhibition at baseline. (top) Average traces at 1.4 x Ethreshold. (bottom) Inhibitory charge plotted as a function of stimulus intensity above Ethreshold. Mixed effects analysis with Dunnett’s Multiple Comparisons, ** p < 0.01, * p< 0.05, n = 21,19, 18,16 cells respectively, 4-6 mice per group. C. Deprivation weakens L4-L2/3 feedforward excitation in wildtype but not Tsc2+/-. (top) Average traces at 1.4 x Ethreshold. (bottom) Excitatory charge plotted as a function of stimulus intensity above Ethreshold. Mixed effects analysis, with Dunnett’s Multiple Comparisons, ** p < 0.01, * p< 0.05, n = 21,19, 18,16 cells respectively, 4-6 mice per group. D. E-I ratio (E/E+I) is elevated in Tsc2+/- sham at 1.2 x Ethreshold and there’s a strong, but nonsignificant trend to increased E-I in WT deprived. Mixed effects analysis, with Dunnett’s Multiple Comparisons, ** p < 0.01, * p< 0.05, n = 21,19, 18,16 cells respectively, 4-6 mice per group. a. Lines with asterisks have symbols on each end to indicate whether comparison is between sham (circle) or deprived (triangle) or both (circle and triangle).

We made miniature postsynaptic current recordings from PV cells to confirm ^20^ that 1-day deprivation does not alter synaptic input onto L2/3 PV cells (**Figure S6**). These experiments used the S5E2 enhancer virus to target PV cells. No differences in miniature excitatory currents (mEPSCs) were observed between sham and deprived mice. In miniature inhibitory (mIPSC) recordings, we observed a small but significant decrease in inter-event interval in PV cells after deprivation, but no change in mIPSC amplitude. These results indicate that brief deprivation does not drive major synaptic changes onto PV cells, suggesting that changes in feedforward inhibition primarily arise from decreased PV excitability.

We hypothesized that in *Tsc2^+/-^* mice, the absence of PV intrinsic plasticity would prevent deprivation-induced weakening of feedforward inhibition in PYR cells. *Tsc2^+/-^*sham mice had less feedforward inhibition than *WT-C57* sham mice (**Figure 6B**), consistent with previous reports^39,40^ and with the reduced PV intrinsic excitability in *Tsc2^+/-^*mice (**Figure 1E**). Feedforward excitation was also reduced at 1.2 x Eθ, but to a smaller degree than inhibition, so that E-I ratio, calculated as E/(E+I), was elevated relative to *WT-C57* (**Figure 6C-D**). Deprivation-induced weakening of feedforward inhibition did not occur in *Tsc2^+/-^* mice, as was weakening of feedforward excitation (**Figure 6B,C**). Thus, the cellular-level deficit in PV intrinsic plasticity was associated with a loss in circuit-level adjustment of feedforward inhibition.

## Discussion

These results show a profound and convergent disruption of PV homeostatic plasticity in two mouse models of autism. In wild-type mice, PV interneurons show bidirectional changes in intrinsic excitability—increasing with sensory use and decreasing with deprivation—strongly supporting their role in homeostatically stabilizing mean firing rate in pyramidal cells. We found that this plasticity relies on activity-dependent regulation of the ER81-*Kcna1* pathway in PV cells, altering expression of Kv1.1 to control PV intrinsic excitability. *Kcna1* gene expression was upregulated within 1.5 hours of deprivation, demonstrating that PV cells rapidly sense and respond to network activity perturbations, faster than other homeostatic mechanisms including synaptic scaling of excitatory synapses ^46^. Deprivation-induced weakening of PV intrinsic excitability was accompanied at the circuit level by reduced feedforward L4-L2/3 inhibition in pyramidal cells, which was lost in *Tsc2^+/-^* mice. This loss of PV homeostasis in *Tsc2^+/-^* and *Fmr1^-/-^* mice at the molecular, cell physiological, and functional circuit levels may contribute to sensory processing dysfunction by maladaptive adjustment or destabilization of circuit function.

### ER81-Kv1.1 pathway for activity-dependent plasticity of PV intrinsic excitability

Sensory experience-dependent regulation of PV intrinsic excitability involved ER81 regulation of *Kcna1* gene expression, which modulated somatic Kv1.1 protein levels in PV cells, and Kv1.1 at the AIS. This pathway was previously shown to mediate PV intrinsic plasticity in response to strong genetic manipulations of PV activity^23^, and the current findings show for the first time its engagement in physiologically relevant sensory-driven plasticity. Kv1.1 powerfully regulates PV spike threshold, spike latency, and overall excitability^21^ and is a common site of regulation for intrinsic plasticity across cell types. This includes for homeostatic plasticity of intrinsic excitability in hippocampal CA3 and CA1 pyramidal cells ^47,48^ and cochlear nucleus neurons^49^, as well as during compensation to gene deletion in several neuron types^50,51^. Kv1.1 expression is transcriptionally regulated by ER81, which directly binds the *Kcna1* promoter to drive gene expression^52^. Other pathways such as neuregulin-ErbB4 and mTOR have also been shown to regulate Kv1 currents in PV cells, and may interact with ER81 to fine-tune PV plasticity^53–55^.

Transcription factor ER81 is likely regulated at the translational level, as proposed previously^23^, to support the observed fast activity-dependent regulation of *Kcna1* that is required for rapid homeostasis by PV cells. Fast activity-dependent gene transcription is well-characterized in the context of immediate early genes such as *Arc* and *Fos*^56–58^. In other cell types, *Kcna1* is rapidly synthesized in dendrites under the control of synaptic activity and mTOR^59^, and can be rapidly removed via selective miRNA-mediated degradation, which may enable rapid bidirectional control of excitability^60^. ER81-based transcriptional activation could contribute to this balance, rapidly adjusting *Kcna1* and other genes expression to regulate intrinsic and synaptic properties of neurons. How ER81 senses activity is unknown, but it is intriguing to consider whether ER81 might act as a synapto-nuclear signal in PV cells, linking changes in synaptic activity to downstream gene transcription.

### Uncoupling of activity-dependent gene regulation in autism

Our results strongly support the hypothesis that, in autism, changes in sensory input are not properly conveyed to the transcriptional or translational machinery, resulting in a loss of the molecular homeostatic response. Dysregulation in activity-dependent gene expression is one of the strongest instances of molecular-level convergence across mouse models of autism ^61,62^. Our results suggest that in *Fmr1^-/-^* and *Tsc2^+/-^* mice, activity is decoupled from ER81 synthesis, disrupting downstream pathways that regulate PV excitability, including Kv1.1 upregulation. This decoupling may occur through different mechanisms—for example in *Fmr1* mice the loss of the RNA-binding function of FMRP could prevent the activity-dependent de-repression of *Kcna1* mRNA. In support of this idea, *Kcna1* was identified as a ‘high-binding’ target of FMRP in hippocampal CA1^63^. In *Tsc2* mice the loss of the Tsc2 brake on mTOR activation might obstruct changes in activity from influencing the protein synthesis machinery^25,36,37^, thus impairing activity-dependent regulation of Kv1.1 or ER81.

### Loss of rapid PV homeostasis in autism

Previous work has shown that homeostatic plasticity of synaptic scaling and intrinsic plasticity of excitatory synapses is impaired in *MeCP2^-/-^*^5–7^, *Fmr1^-/-^* ^8,64^, *Chd8^+/-^* ^9^, and *Shank3^-/-^* ^10,11^ models of autism. Our finding that PV circuit homeostasis is also impaired suggests a broad deficit in activity-dependent homeostatic mechanisms. Impaired homeostasis could leave networks vulnerable to perturbations ^4^, could drive networks to develop the wrong set point of activity, impair information flow, or degrade neural tuning^3^. Because of the critical role of PV cells in regulating Hebbian plasticity, impairment of PV homeostasis may promote maladaptive plasticity in response to sensory use, compromising sensory processing and perception.

We verified that the inability of deprivation to weaken PV intrinsic excitability in *Fmr1^-/-^* and *Tsc2^+/-^* mice represents a true loss of PV homeostasis and not a ‘floor effect’ in which the process of homeostatic downshifting of PV excitability is intact, but already maximally activated in whisker-intact animals. First, ER81 and Kv1.1 levels, which normally increase with deprivation, are normal or reduced, not increased, in whisker-intact *Fmr1^-/-^* and *Tsc2^+/-^*mice (**Fig. 3**), suggesting that the homeostatic downshifting mechanism is not already activated or saturated. Second, if homeostatic downshifting were partially or maximally engaged at baseline, whisker enrichment should increase (upshift) PV excitability in *Fmr1^-/-^* and *Tsc2^+/-^* mice more than wild type mice. We tested this in *Tsc2^+/-^*mice and upshifting was not increased but abolished (**Fig. 5**). Thus, PV cells exhibit a fundamental loss of bidirectional intrinsic homeostasis. We hypothesize that this arises from disruptions in the FMRP- and mTOR-dependent molecular signaling pathways that mediate activity-dependent gene regulation. These disruptions may include not just ER81 and Kv1.1 but other pathways as well.

Contribution of PV intrinsic excitability and impaired homeostasis to PV hypofunction in autism PV hypofunction, identified by either reduction in PV cell numbers, PV spiking, or net PV inhibitory output, has been observed in many autism mouse models including *Cntnap2^-/-^*, *Shank3^-/-^*, *Scn1a^+/-^,* and *Chd8^+/-^,* and in postmortem human brain tissue ^12,14,15,65^. Some autism gene mutations directly reduce PV cell numbers, for example by impairing cell migration during development^66^. Other genes affect synaptic excitation onto PV cells, including *Shank3* which regulates development of excitatory synapses onto PV cells^67,68^. Some autism risk genes may directly impair intrinsic excitability of PV cells, including *Cntnap2* which is essential for clustering potassium channels in the AIS and axons^66,69,70^, and *Scn1a* which encodes the primary sodium channel necessary for PV cell spiking ^71,72^.

Our findings demonstrate that PV intrinsic excitability is also reduced in *Fmr1^-/-^*and *Tsc2^+/-^* models of autism, evident either as a reduction in the F-I curve or as an elevation in PV spike threshold (**Fig. 1).** Because ER81 and Kv1.1 levels were not elevated at baseline in these mice (**Fig. 3**), this alteration in excitability is not likely due to specific disruption in Kv1.1 expression but must involve other ion channels. In *Fmr1^-/-^* mice, this may reflect translational dysregulation of downstream ion channel genes (e.g., Kv3.1 and Kv4.2) or modulators, and/or reduced FMRP binding to BK, SLACK, HCN and Cav2.2 channels, which can all contribute to altered excitability^73,74^. In *Tsc2^+/-^* mice, disrupted mTOR signaling may also alter expression of ion channels to alter excitability^39^. We hypothesize that impaired PV intrinsic excitability contributes to PV hypofunction for other autism risk genes that converge on common pathways regulating ion channel expression, potentially including *Chd8* and *Arid1b*^75,76^. Reduced PV intrinsic excitability may act as an additional contributing factor to reduce PV cell numbers, because PV activity regulates PV cell apoptosis during development^77,78^.

Whether impaired PV homeostasis is itself a cause of PV hypofunction in autism is unknown. Developmental strengthening of synapses and intrinsic excitability in PV circuits may be driven homeostatically by developmental increases in excitatory network activity. If PV homeostasis is impaired, such developmental strengthening of PV circuits may not occur.

### Relationship between PV homeostasis and E-I ratio in autism

A prior paper showed that 4 autism mouse models, including *Fmr1^-/-^*and *Tsc2^+/-^*, exhibited a coordinated reduction in feedforward excitation and inhibition in S1 that elevated E-I ratio but was quantitatively balanced to stabilize, not increase, peak postsynaptic depolarization and firing rate in L2/3 PYR neurons. This initially suggested that cellular homeostasis mechanisms were intact in autism, and that PV hypofunction could arise as an active homeostatic compensation to a primary excitatory circuit deficit ^40^. However, our new data add an additional piece of the puzzle—multiple forms of homeostasis coexist within circuits and may compensate when other forms fail ^40,79^. Thus, while rapid PV homeostasis is lost in *Fmr1^-/-^* and *Tsc2^+/-^* models, slower forms of homeostasis may still operate to balance E and I to stabilize mean PSPs and mean firing rate in autism during development or on longer timescales (days to weeks). If circuits in autism lack a subset of homeostatic mechanisms, they may be able to adapt within a range, or on slow timescales, but when challenged with dynamic perturbations to activity, may lose their resilience. This could have dramatic downstream consequences by facilitating maladaptive Hebbian plasticity at feedforward synapses, potentially driving changes in thalamocortical connectivity, leading to cognitive inflexibility, insistence on sameness and stereotyped behaviors. Future work exploring the dynamics of homeostatic compensation across excitatory and inhibitory networks is necessary to clarify how autism circuits adapt or fail under stress. Understanding the balance between slow and rapid homeostatic processes could reveal novel approaches to enhance circuit resilience.

## Concluding remarks

Our findings establish impaired PV circuit homeostasis as a shared circuit defect in at least two genetic forms of autism, mediated by decoupling the ER81-Kv1.1 pathway from control by neural activity. These results provide critical insight into the molecular and circuit-level mechanisms underlying sensory processing deficits in autism and point to potential targets for therapeutic intervention. Future studies exploring PV homeostasis across other autism models and its impact on behavior will be essential to further elucidate its role in ASD pathophysiology.

## Supporting information

Supplemental Table 1- Statistical Analysis

## Acknowledgements

We are grateful to all Feldman Lab members for their feedback and discussions. We thank the CRL Molecular Imaging Center (MIC) at Berkeley (**RRID:SCR_017852)** for their assistance with imaging troubleshooting and optimization. The microscopes used are supported by the Helen Wills Neuroscience Institute. This work was funded by 1F32 NS126310 (HRM), NSF Graduate Research Fellowship (AMN), R01 NS105333 (DEF) and SFARI Investigator Award grant (DEF).

## Author Contributions

Conceptualization, H.R.M. and D.E.F.; Methodology, H.R.M., H.C.W., A.M.N. and D.E.F.; Investigation, H.R.M., A.M.N., H.C.W., S.A.Y., K.W.W., F.E.B.; Software, H.R.M., S.L.; Writing –Original Draft, H.R.M. and D.E.F.; Writing – Review & Editing, H.R.M., A.M.N., and D.E.F.; Funding Acquisition, H.R.M. and D.E.F.; Supervision, H.R.M. and D.E.F.

## Methods

### Experimental Model

Procedures were approved by the University of California, Berkeley Animal Care and Use Committee. PV-IRES-Cre mice (Strain #008069, The Jackson Laboratory) were crossed with Cre-dependent TdTomato reporter (Ai14) mice (The Jackson Laboratory; Strain#007914) to generate PV-Cre;tdTomato offspring. C57-Fmr1-/- mice (Strain# 003025) and Tsc2+/- mice (Strain #004686) were obtained from Jackson Laboratory and bred in house. WT-C57 mice were obtained from Charles River. Mice were housed as litters in standard cages. Littermates of either sex were randomly assigned to experimental groups. Males and females were used for experiments with equal frequency. No sex-specific differences were noted. Deprived/Enriched mice and sham mice were littermates, and were recorded interleaved either on the same day or on alternate days. WT mice are a mix of Tsc2+/+ and C57 mice in separate litters, and were recorded interleaved with Tsc2+/- littermates or Fmr1-/- respectively. For whisker deprivation, the right D-row whiskers (D1–D6 and gamma) were plucked under transient isoflurane anesthesia, 24 ±2 h before slice preparation.

Sham-plucked littermates underwent anesthesia but not plucking. For whisker enrichment, all whiskers except the right D-row (D1-D6 and gamma) were plucked under transient isoflurane anesthesia. Sham-plucked littermates underwent anesthesia but not plucking. Immediately following recovery from anesthesia, sham-plucked animals were weaned into standard cages (2 mice per cage) and D-row spared animals were weaned into enriched environment cages (3 mice per cage) containing a plastic hut (Kaytee Small Animal Igloo Hideout), plastic tube (Kaytee CritterTrail), and 3- 4 other textured wooden or plastic toys. Animals remained in these housing conditions for 1-3 days (24-72 ±2 h) before slice preparation.

### Method Details

#### Neonatal Viral Injection

Mice were injected with AAV9-S5E2-dTom-nlsdTom (www.addgene.org/135630/) or AAV9**-**S5E2- GCaMP6f (ttps://www.addgene.org/135632/). The dTom virus used for Figure 1 was made by Vector Biolabs, whereas the enrichment experiments used virus from Addgene. Mice age P2-P4 were anesthetized on ice for 2-3 minutes and then rapidly injected using a Nanoject. 414 nl were injected at one site in the left hemisphere (coordinates from lambda: AP: +1.60 mm, ML: +1.75 mm, ∼0.3 mm DV) for PV intrinsic excitability recordings. Mice were returned to the mother with their littermates and allowed to recover for 2-3 weeks

#### Slice preparation

Postnatal day 18 (P18) to P22 mice of either sex were anesthetized with isoflurane and decapitated. Brain slices were prepared using a Leica VT1200S vibratome in chilled, oxygenated, low-sodium, low-calcium Ringer’s solution (in mM: 85 NaCl, 75 sucrose, 25 D-()-glucose, 4 MgSO4, 2.5 KCl, 1.25 Na2HPO4, 0.5 ascorbic acid, 25 NaHCO3, and 0.5 CaCl2, 320 mOsm). Cortical slices (350 µm) were cut from the left hemisphere in the “across-row” plane, oriented 50° toward coronal from the midsagittal plane and 35° from vertical. Using this plane, each slice contains one column from each whisker row A–E, and within-column circuits are largely preserved^80,81^. Slices were transferred to standard Ringer’s solution (in mM: 119 NaCl, 26.2 NaHCO3, 11 D-()-glucose, 1.3 MgSO4, 2.5 KCl, 1 NaH2PO4, 2.5 CaCl2, 300 mOsm) for 30 min at 30°C and then were kept at room temperature until recording (0.5–6 h).

#### Slice electrophysiology

Recordings were made at 30 –31°C in standard Ringer’s solution. Barrel columns were identified by transillumination and visually guided patching was performed using infrared differential interference contrast optics at 40X. L2/3 PYR cells were identified by soma shape, and were located 100 –240 µm below the L1–L2 boundary, within 100 µm tangentially of column center. PV neurons were identified by tdTomato fluorescence (530 –550 nm bandpass excitation, 575– 625 nm emission; Moment CMOS camera, Teledyne Photometrics). All recordings were made in L2/3 of D-row barrel columns, with the NMDA receptor antagonist 50 M D-AP5 in the bath. Whole-cell recording was performed with 3–5 MΩ pipettes using a Multiclamp 700B Amplifier (Molecular Devices) with 2 kHz low-pass filtering and 7–10 kHz digitization. L4-evoked synaptic responses were elicited using 200 µs current pulses delivered via a bipolar stimulating electrode (115 m tip spacing, FHC) placed in the center of the D barrel in L4. The interstimulus interval was 10 s. For input– output curves, the EPSC threshold (E) was defined as the minimal stimulation current that evoked an EPSC in five consecutive sweeps. For L2/3 PYR cells, the E was determined individually for each cell. Voltage-clamp recordings of synaptic currents used cesium gluconate internal solution (in mM: 108 D-gluconic acid, 108 CsOH, 20 HEPES, 5 tetraethylammonium (TEA)-Cl, 2.8 NaCl, 0.4 EGTA, 4 MgATP, 0.3 NaGTP, 5 BAPTA, and 5 QX-314 bromide, pH 7.2, 295 mOsm). The holding potential (Vhold) was corrected for the liquid junction potential (12 mV). Series resistance (Rseries) was monitored in each sweep and was compensated by 40 – 80%. Cells whose input resistance (Rinput) changed >30% throughout recording were excluded from analysis. PYR cells with a resting membrane potential (Vrest) that was more depolarized than -60 mV were discarded. Spontaneous miniature EPSC and IPSCs were recorded in voltage clamp at Vhold -72mV or 0 mV with 50 µM d- AP5 and 0.1 µM tetrodotoxin (TTX) in the bath. Rseries compensation was not used for miniature recordings. Events were detected using a deconvolution algorithm based on Pernía-Andrade et al. (2012), with a noise threshold of 3.5x. All events larger than ± 2 pA that were separated by 1 ms were analyzed and subsequently screened manually. The detection threshold was 5 pA. A minimum of 300 events was analyzed per cell.

Current-clamp recordings were performed using K gluconate internal solution (mM: 116 K gluconate, 20 HEPES, 6 KCl, 2 NaCl, 0.5 EGTA, 4 MgATP, 0.3 NaGTP, and 5 Na2phosphocreatine, pH 7.2, 290 mOsm). Series resistance artifacts were corrected by bridge balance. PV cell intrinsic excitability was recorded in whole-cell current clamp mode. Vrest was measured immediately after break-in.

Input resistance was calculated as the slope of the linear fit of steady-state Vm during -50, 0, and 50 pA current steps. Rheobase was defined as the minimum current injection (500 ms) that elicited one or more spikes on 3 out of 5 consecutive sweeps. The firing– current (F–I) relationship was measured using increasing currents above rheobase. Spike threshold was defined as the Vm at which the second derivative of Vm was >6 SDs above the prestimulus period. Spike latency was defined as the time to spike threshold. Cells with a spike threshold of less than -20 mV, indicative of an undercompensated bridge, were excluded from analysis. Cells with an input resistance less than 50 MΩ were excluded from analysis.

#### Immunohistochemistry

Mice were anesthetized and then perfused with 2% PFA. Brains were then fixed in 4% PFA for 2 hours and then transferred to 30% (w/v) sucrose solution and stored at 4 °C. Fixing overnight resulted in loss of PV immuno signal. 50-μm across-row sections were cut using a freezing microtome. Free-floating sections were initially blocked with a solution containing 2% Bovine Serum Albumin(BSA), 0.125% Triton X-100, and 10% Normal Goat Serum (GCS) during 1 h at room temperature. Incubation with primary antibodies was performed overnight at 4 °C. As primary antibodies, PV antibody (rabbit anti-Parvalbumin, clone 4G3, EMD Millipore Corp., USA ; 1:200), PV antibody (mouse anti-Parvalbumin ; 1:200), the Kv1.1 antibody (mouse monoclonal anti-Potassium Channel kv1.1 channel, clone K36/15, EMD Millipore Corp., USA ; 1:250 dilution), the AnkG antibody (mouse anti-Ankyrin monoclonal, clone N106/36, EMD Millipore Corp., USA; 1:1000), the VGLUT2 antibody (guinea pig polyclonal, AB2251-1, EMD Millipore Corp., USA; 1:1000), the RPS6 antibody (rabbit anti-P-S6 Ribosomal Protein S240/244, D68F8, Cell Signaling Technology; 1:800), the ER-81 antibody (rabbit anti-ER-81, MBS460434, MyBioSource; 1:200), and the PGC-1a antibody (mouse anti-PGC-1a, sc-518025, Santa Cruz Biotechnology; 1:200 dilution), were used. Next, sections were washed three times with 0.1M PBS, for 5 min each, before incubating with secondary antibodies. The following secondary antibodies were used: anti-rabbit Alexa488-conjugated antibody (Life Technologies Corporation, Eugene, OR; 1:500), anti-mouse Alexa647-conjugated antibody (Life Technologies Corporation, Eugene, OR; 1:250), anti-guinea pig Alexa647-conjugated antibody (Life Technologies Corporation, Eugene, OR; 1:400 dilution), anti-rabbit Alexa568-conjugated antibody (Life Technologies Corporation, Eugene, OR; 1:250), anti-mouse Alexa568-conjugated antibody (Life Technologies Corporation, Eugene, OR; 1:500), anti-guinea pig Alexa568-conjugated antibody (Life Technologies Corporation, Eugene, OR; 1:400), and anti-mouse Alexa488-conjugated antibody (Life Technologies Corporation, Eugene, OR; 1:500). Sections were incubated with secondary antibodies overnight at 4 °C. Finally, all sections were washed three times with 0.1M PBS and stained with DAPI (Life Technologies Corporation, Eugene, OR; 1:1000 dilution), for 20 min in 0.1M PBS. Stained sections were imaged using a Zeiss LSM 880 NLO laser scanning confocal microscope. Images were acquired in Layer l2/3 of the D column using staining for vGlut to delineate the barrels and/or PV staining and DAPI. Images were acquired at 20X for Figure 3 and Figure 5 in D column of Layer ⅔. For analysis of somatic signal inside PV-positive cells, a custom ImageJ (FIJI) macro was written to automatically obtain PV cell regions of interest (ROI) using Threshold and Analyze Particles functions in the PV channel. This ROI mask was then applied to the other channels (Kv1.1, ER81 etc) to measure fluorescence intensity. Fluorescence intensity inside ∼ 10- 30 PV cells was averaged for each slice. For measurement of axon initial segments in Figure 2, images were acquired using high-resolution Airyscan at 63X and 1.8 zoom. PV cells in L2/3 of the D column that had a clearly associated AIS (as seen with AnkG marker) were imaged. Images underwent Airyscan post-processing prior to analysis. During analysis, the AIS associated with the PV cell was re-identified by a trained technician by scrolling through the z-stack. For AIS measurements, the z-stack was max-projected. The length of the AIS was measured on the max-projected image by drawing a line in the center of the AIS from soma to end. The Kv1.1 signal was measured by drawing an ROI around the AnkG signal of the AIS and measuring Kv1.1 inside the ROI. The distance of the AIS from the soma was measured as a straight line from the edge of the PV somatic signal to the start of the AIS. Any AIS’s that overlapped with Kv1.1-positive somas of other cells in the tissue were excluded.

All imaging and analysis was performed blind to condition. Each imaging dataset is the result of at least two technical replicates (batches) and a minimum of 3 mice per group. All imaging and analysis parameters including but not limited to laser power, gain, pixel size, and threshold settings were kept consistent across each batch. All batches contained all relevant experimental groups so that comparisons between groups were well-controlled.

### HCR-FISH

HCR-FISH experiments were conducted on thick ‘across-row’ slices (100 or 350 µm). Slices were cleared with 8% SDS for 2 hrs at RT, then washed with 2X SSC to remove all SDS. Probes B1-PV (4 nM) and B5-Kcna1 (16 nM) (Molecular Instruments) were used. Slices were incubated with probes for 24 hours at 37C and 24 hours at RT before proceeding to amplification. B1-488 and B5- 647 were used for amplification. Sections were imaged on the Zeiss LSM 880 NLO laser scanning confocal microscope using the same laser power and gain for each batch.

Sham and deprived slices were run in the same batch and analyzed with the same parameters using a custom application of the big-FISH package (https://github.com/fish-quant/big-fish). Settings for analysis including threshold and spot size were kept consistent for each batch. The qHCR-FISH images contained three channels: Kcna1 mRNA (qHCR-FISH probe), cell (PV cell mRNA), and nuclear (DAPI). The pipeline comprised three major components: spot detection, cell and nuclear segmentation, and integrative analysis. For spot detection, we applied Gaussian and log filters to preprocess the images, adjusting the preprocessing methods for each batch as needed. Bright spots corresponding to Kcna1 RNA were identified using the spot detection functions in Big-FISH. Additionally, we detected and decomposed condensed regions to better quantify RNA levels in areas where multiple RNA molecules overlapped. For cell and nuclear segmentation, binary masks were created using ImageJ to segment PV cells and nuclei. These masks were subsequently converted into labeled images, assigning unique identifiers to individual cells. Finally, the results from spot detection and segmentation were integrated to quantify Kcna1 mRNA levels in each PV cell. The integrative analysis results were normalized to generate statistics for comparing Kcna1 mRNA levels across different conditions. Our code and documentation for the pipeline has been uploaded in https://github.com/Luuuuux/FISH-quant.

### Quantification and Statistical Analysish

Statistical analysis was performed in Matlab or Graphpad Prism. P-values, statistical test used, and ‘n’ are reported in the Figure Legends. Table S1 contains full details of statistical analysis for each figure panel. All reported effects were observed across at least three independent litters.

Experiments were repeated at least three times (technical replicates). At least 3 mice per group were used in every experiment (biological replicates). For each measurement type, data were tested for Gaussian distribution before use of parametric statistics. Non-Gaussian data were evaluated using nonparametric statistics

Figure S1 (related to Figure 1). Passive properties and spike waveform are largely unaffected by deprivation or genotype.

A. Input resistance in PV cells across genotypes and deprivation conditions is not changed. Two-way ANOVA, p>0.05
B. Rheobase in PV cells across genotypes and deprivation conditions is not changed. Two-way ANOVA, p>0.05
C. Vrest in PV cells across genotypes and deprivation conditions is not changed. Two-way ANOVA, p>0.05
D. Peak action potential amplitude in PV cells across genotypes and deprivation conditions is not changed. Spike properties measured in all spikes at 0, 10, 25, and 50 pA above rheobase.Two-way ANOVA, p>0.05
E. Full width half height of action potential is increased in *Tsc2*+/- sham. Two-way ANOVA with Tukey’s test for Multiple Comparisons, Genotype: F(2,86) = 4.333, p = 0.0161
F. Fast afterhyperpolarization depth is not changed. Two-way ANOVA, p>0.05
G. (n = 22,19, 22,15, 13,14 cells, 5-7 mice per group)

Figure S2 (Related to Figure 4). Additional quantification of Kcna1 mRNA in PV cells

A. Scatterplots showing raw mRNA counts per cell at different timepoints after deprivation (same data at Figure 4C). Line and error bars represent mean + SEM. Symbols represent slices. 1.5 hr: Unpaired t-test, p = 0.0169
B. Scatterplots showing raw mRNA counts per nucleus at different timepoints after deprivation (same data at Figure 4C). Line and error bars represent mean + SEM. Symbols represent slices. 1.5 hr: Unpaired t-test, p = 0.0449

Figure S3 (related to Figure 5). Passive properties and spike waveform in sparing plus enrichment experiments.

A. Immediate early gene c-fos, a marker of neuronal activation, is enhanced in L2/3 of the D column (white arrow) following whisker enrichment paradigm (EE+ D-row sparing).
B. Input resistance in PV cells across genotypes and experience conditions is not changed. Two-way ANOVA, p>0.05
C. Rheobase in PV cells is significantly different between WT-C57 and Tsc2+/- mice. Two-way ANOVA with Tukey’s test for Multiple Comparisons, Genotype: F(1,78) = 4.951, p = 0.029.
D. Vrest in PV cells is significantly different between WT-C57 and Tsc2+/- mice. Two-way ANOVA with Tukey’s test for Multiple Comparisons, Genotype: F(1,78) = 4.027, p = 0.048.
E. Peak action potential amplitude in PV cells across genotypes and experience conditions is not changed. Spike properties are measured in all spikes at 0, 10, 25, and 50 pA above rheobase. Two-way ANOVA, p>0.05
F. Full width half height of action potential across genotypes and experience conditions is not changed. Two-way ANOVA, p>0.05
G. Fast afterhyperpolarization Two-way ANOVA with Tukey’s test for Multiple Comparisons, Genotype: F(1,78) = 6.419, p = 0.013.

Figure S4 (related to Figure 6). mEPSCs, mIPSCs, and feedforward synaptic response properties in PV cells

A. Representative traces from mini EPSCs recorded in fluorescent PV cells from PV-tdTomato mice (C57 background).
B. Average mEPSC waveform from all cells from sham and deprived mice.
C. mEPSC amplitude is not altered by deprivation. Mann-Whitney test, p>0.05
D. mEPSC frequency is not altered by deprivation. Mann-Whitney test, p>0.05
E. Representative traces from mini IPSCs recorded in fluorescent PV cells from PV-tdTomato mice (C57 background).
F. Average mIPSC waveform from all cells from sham and deprived mice.
G. mIPSC amplitude is not altered by deprivation. Mann-Whitney test, p>0.05
H. mIPSC frequency is significantly increased with deprivation. Mann-Whitney test, p<0.05, n = 12, 13 cells from 4 and 5 mice respectively
I. Rinput is not changed in voltage clamp recordings from wildtype and deprived WT and Tsc2+/- mice. Two-Way ANOVA n = 21,19, 18,16 cells respectively, 4-6 mice per group.
J. Age is slightly higher in WT deprived mice. Two-Way ANOVA with Tukey’s test for Multiple Comparisons, * p < 0.05. n = 21,19, 18,16 cells respectively, 4-6 mice per group.
K. E threshold is higher in Tsc2+/- mice than wildtype. Two-Way ANOVA with Tukey’s test for Multiple Comparisons, Genotype effect, **p = 0.0023 n = 21,19, 18,16 cells respectively, 4-6 mice per group.
L. Scatterplot showing distribution of all cells for inhibitory (left) and excitatory (right) charge at 1.2 X Ethreshold. Corresponds to Figure 5B, C. Two-Way ANOVA with Tukey’s test for Multiple Comparisons, *p <0.05, **p < 0.01, ***p<0.001, n = 9,8,11,7 cells, 3-5 animals

