## Supplemental Figures for "Physiological and molecular impairment of PV circuit homeostasis in mouse models of autism"

Figure S1

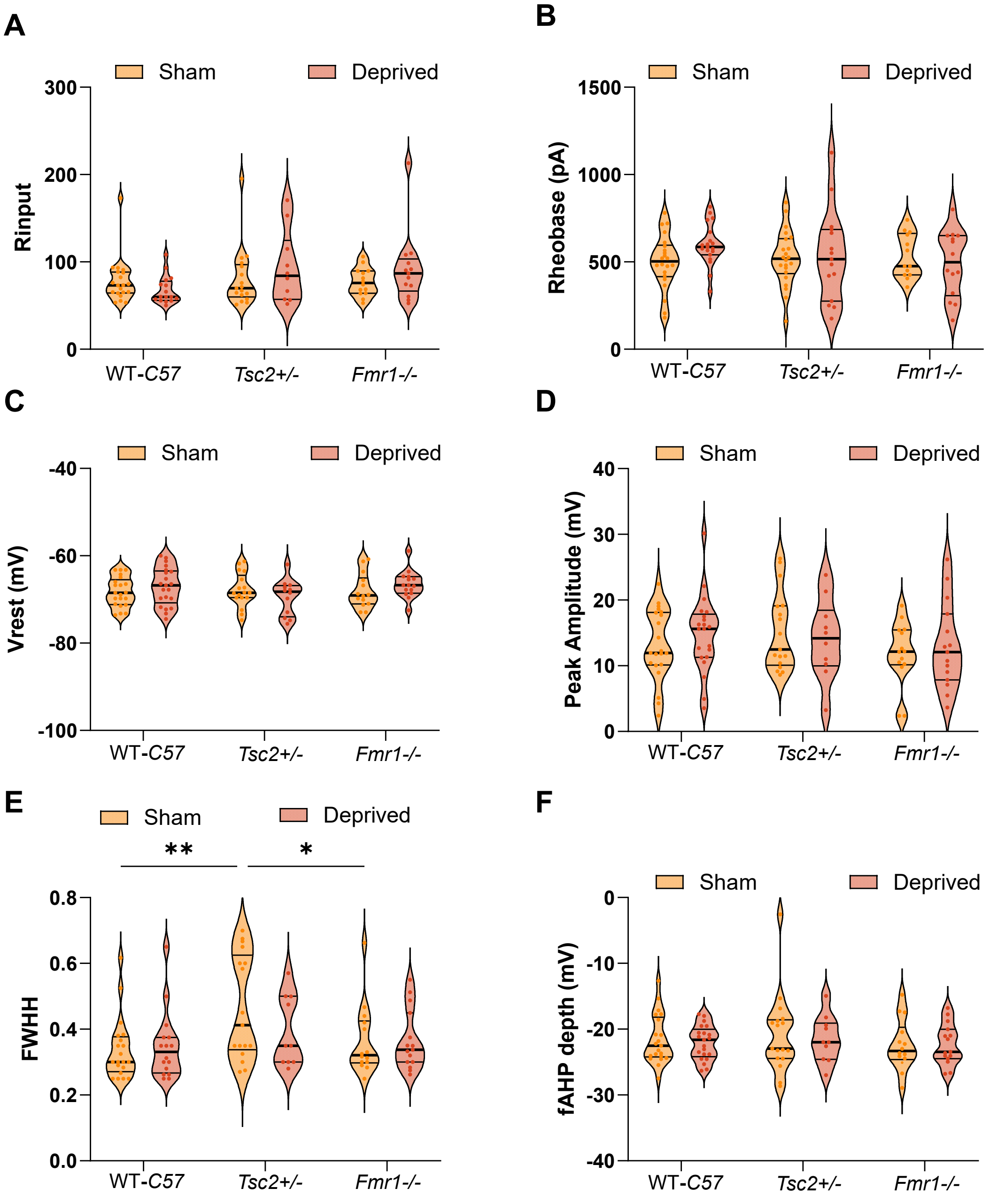


Figure S1 (related to Figure 1). Passive properties and spike waveform are largely unaffected by deprivation or genotype.

Figure S2

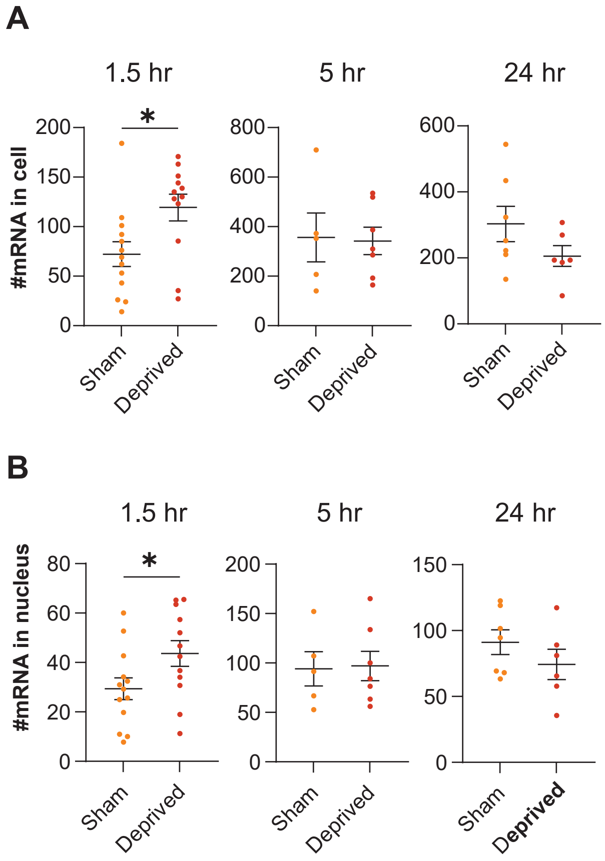


Figure S2 (Related to Figure 4). Additional quantification of Kcna1 mRNA in PV cells

Figure S3

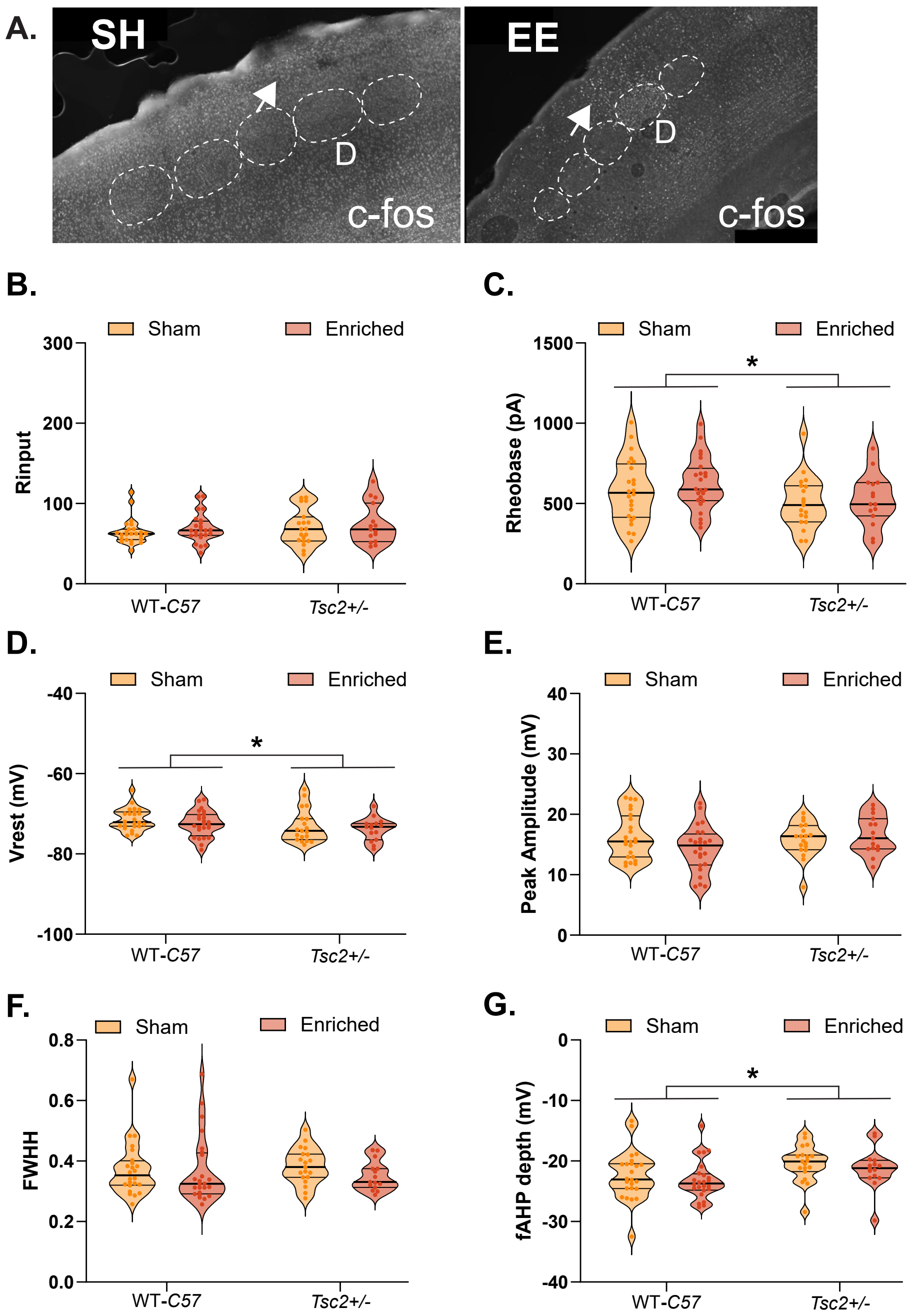


Figure S3 (related to Figure 5). Passive properties and spike waveform in sparing plus enrichment experiments.

Figure S4

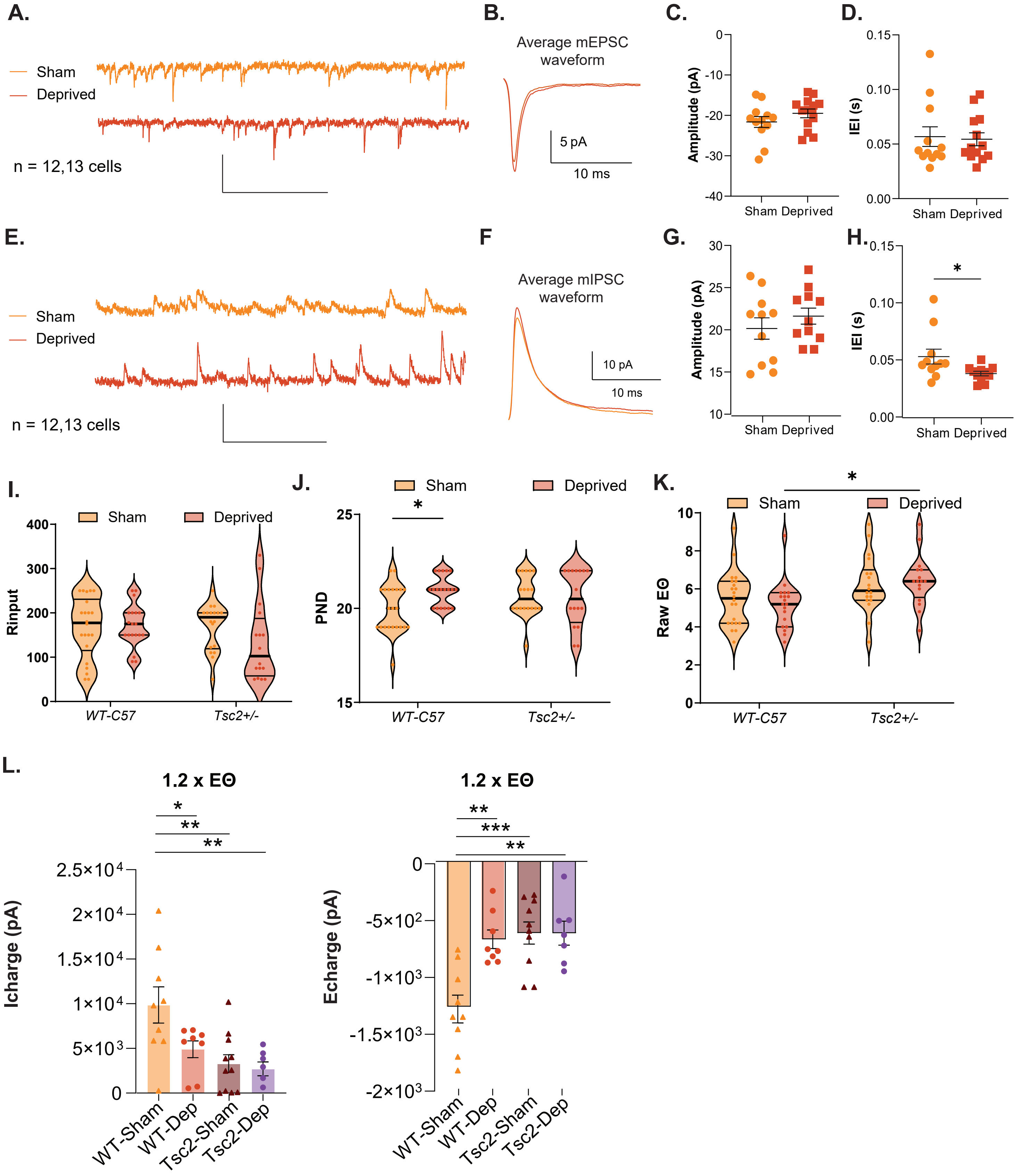


Figure S4 (related to Figure 6). mEPSCs, mIPSCs, and feedforward synaptic response properties in PV cells
